# Cockayne syndrome proteins CSA and CSB maintain mitochondrial homeostasis through NAD^+^ signaling

**DOI:** 10.1101/2020.03.01.972323

**Authors:** Mustafa N. Okur, Evandro F. Fang, Elayne M. Fivenson, Vinod Tiwari, Deborah L. Croteau, Vilhelm A. Bohr

## Abstract

**Background:** Cockayne syndrome (CS) is a rare premature aging disease, most commonly caused by mutations of the genes encoding the CSA or CSB proteins. CS patients display cachectic dwarfism and severe neurological manifestations and have an average life expectancy of 12 years. The CS proteins are involved in transcription and DNA repair, with the latter including transcription-coupled nucleotide excision repair (TC-NER). However, there is also evidence for mitochondrial dysfunction in CS, which likely contributes to the severe premature aging phenotype of this disease. While damaged mitochondria and impaired mitophagy were characterized in mice with CSB deficiency, such changes in the CS nematodes and CS patients are not fully known.

**Results:** Our cross-species transcriptomic analysis in CS postmortem brain tissue, CS mouse and nematode models show that mitochondrial dysfunction is indeed a common feature in CS. Restoration of mitochondrial dysfunction through NAD^+^ supplementation significantly improved lifespan and healthspan in the CS nematodes, highlighting mitochondrial dysfunction as a major driver of the aging features of CS. In cerebellar samples from CS patients, we found molecular signatures of dysfunctional mitochondrial dynamics and impaired mitophagy/autophagy. In primary cells depleted for CSA or CSB, this dysfunction can be corrected with NAD^+^ supplementation.

**Conclusions:** Our study provides support for the interconnection between major causative aging theories, DNA damage accumulation, mitochondrial dysfunction, and compromised mitophagy/autophagy. Together these three agents contribute to an accelerated aging program that can be averted by NAD^+^ supplementation.

## Background

Patients with Cockayne Syndrome (CS) experience cachectic dwarfism and microcephaly, with an average life expectancy of 12 years. CS is considered a segmental aging disease and a model of premature aging. Along with increased genetic damage, mitochondrial dysfunction has been implicated in CS progression; however, the mechanism of this dysfunction remains to be further explored [1-4].

CS is predominantly caused by mutations in the genes encoding either the CSA or CSB proteins. CSA [5, 6] and CSB [7] are DNA damage response proteins that are involved in transcription-coupled nucleotide excision repair (TC-NER), as well as both nuclear and mitochondrial base excision repair (BER) [8]. NER is a versatile DNA repair pathway as it is responsible for the removal of a wide range of DNA adducts. CSA and CSB are also important for transcription recovery after DNA damage [9-11]. Various studies have shown that CSB plays a considerable role in transcription, and likely is a transcription factor [12-15].

Similar to other DNA repair-defective premature aging diseases, such as xeroderma pigmentosum (XP)[16], ataxia-telangiectasia (AT) [17], and Werner syndrome (WS) [18], mitochondrial dysfunction is implicated in many CS phenotypes [3, 4, 19]. The mechanisms of mitochondrial dysfunction underlying CS remain unclear but may involve the impairment of the NAD^+^(nicotinamide adenine dinucleotide, oxidized)-mitophagy axis [19-22]. NAD^+^ is a fundamental molecule in cellular energy metabolism and signaling pathways and is essential for adaptive responses of cells to bioenergetics and oxidative stress; accordingly, there is an age-dependent depletion of cellular NAD^+^, suggesting NAD^+^ depletion as a major driver of both pathological and biological aging [23]. Many molecular mechanisms are involved in the multi-faceted effects of NAD^+^ on longevity and healthspan, including the induction of mitophagy, a cellular self-recognition, engulfment, degradation, and recycling of damaged and superfluous mitochondria [24-26]. In CS, the accumulation of unrepaired DNA damage may lead to mitochondrial dysfunction through increased PARP-1 mediated NAD^+^ consumption [4]. Decreased NAD^+^ levels lead to decreased activity of SIRT-1, an NAD^+^-dependent deacetylase that maintains mitochondrial homeostasis through coordination of elimination of damaged mitochondria via mitophagy and mitochondrial biogenesis via activation of PGC-1α and FOXOs [16]. This phenomenon has also been observed in other DNA repair disorders [16, 17]. Accordingly, rapamycin treatment or dietary restriction, which upregulate mitophagy, has shown promise in CS models [19] [22]. It is therefore examined whether increasing NAD^+^ levels through NAD^+^ precursor supplementation can ameliorate symptoms of mitochondrial dysfunction in CS.

Our cross-species and unbiased transcriptomic analysis on CS postmortem brain tissue, CS mouse cerebellum, and CS *Caenorhabditis elegans* (*C. elegans*) models show Gene Ontology (GO) terms associated with mitochondria are changed in all CS models and normalized with NAD^+^ supplementation in worms and mouse models of CS. We also show that various mitochondrial abnormalities such as increased mitochondrial length and reduced mitochondrial networking are present in CS worms and are restored by NAD^+^ precursor treatment. Lastly, the activity of several key proteins of mitochondrial homeostasis such as AMPK and ULK-1 are impaired in the CS postmortem brain tissues. NAD^+^ supplementation restores the activity of these proteins in primary cell lines deficient in CSA or CSB. This has implications for intervention in CS and possibly aging.

## Results

### Microarray analysis of CS patient cerebellum samples

The loss of CSA and CSB proteins is known to compromise the central nervous system function, with patients exhibiting varying degrees of demyelination, neuronal loss, calcification, gliosis and cerebellar atrophy [1]. To examine changes in gene expression in the cerebellum of CS patients, microarray analysis was conducted using frozen cerebellar samples from CS patients and age-matched controls from the NIH NeuroBioBank’s Brain and Tissue repository at the University of Maryland, Baltimore (Supplementary Table 1). Since the origin of the gene mutations in these patients is unknown, hierarchical clustering was performed on the samples to see if the CS patient-derived samples clustered together (Supplementary Fig. 1). From this analysis, two samples, CS2 and CS4R, clustered more closely with the controls (WT) than the CS patients and were therefore removed from further array analysis. We used gene set enrichment analysis to parse the genes into various GO terms and identified 166 (88%) up-regulated and 22 (12%) down-regulated terms (Supplementary Table 2). A heatmap depicting Z-scores comparing the change between CS patients and controls is shown with the various GO terms sorted into related topics (Figure 1a). Terms associated with mitochondrial function, synapse and neuronal function, immune response, and cellular stress response were the largest groups among all the GO terms. Notably, all of the terms associated with synapse and neuronal function were significantly down-regulated. In contrast, many terms associated with mitochondrial function, immune response, and stress response pathways were significantly up-regulated. Taken together, the signature of changes in CS patient samples relative to controls suggest a proinflammatory, oxidative stress environment with neuronal cells suffering from mitochondrial and synaptic dysregulation (Fig. 1a). Previously, we implicated oxidative stress, persistent DNA damage signaling and NAD^+^ depletion as contributors to mitochondrial dysfunction in mice with CSB deficiency [4, 19]. Our microarray data of human postmortem brain tissues show cross-species consistency of these affected terms and are therefore of great interest in understanding CS pathology.

**Figure 1.**
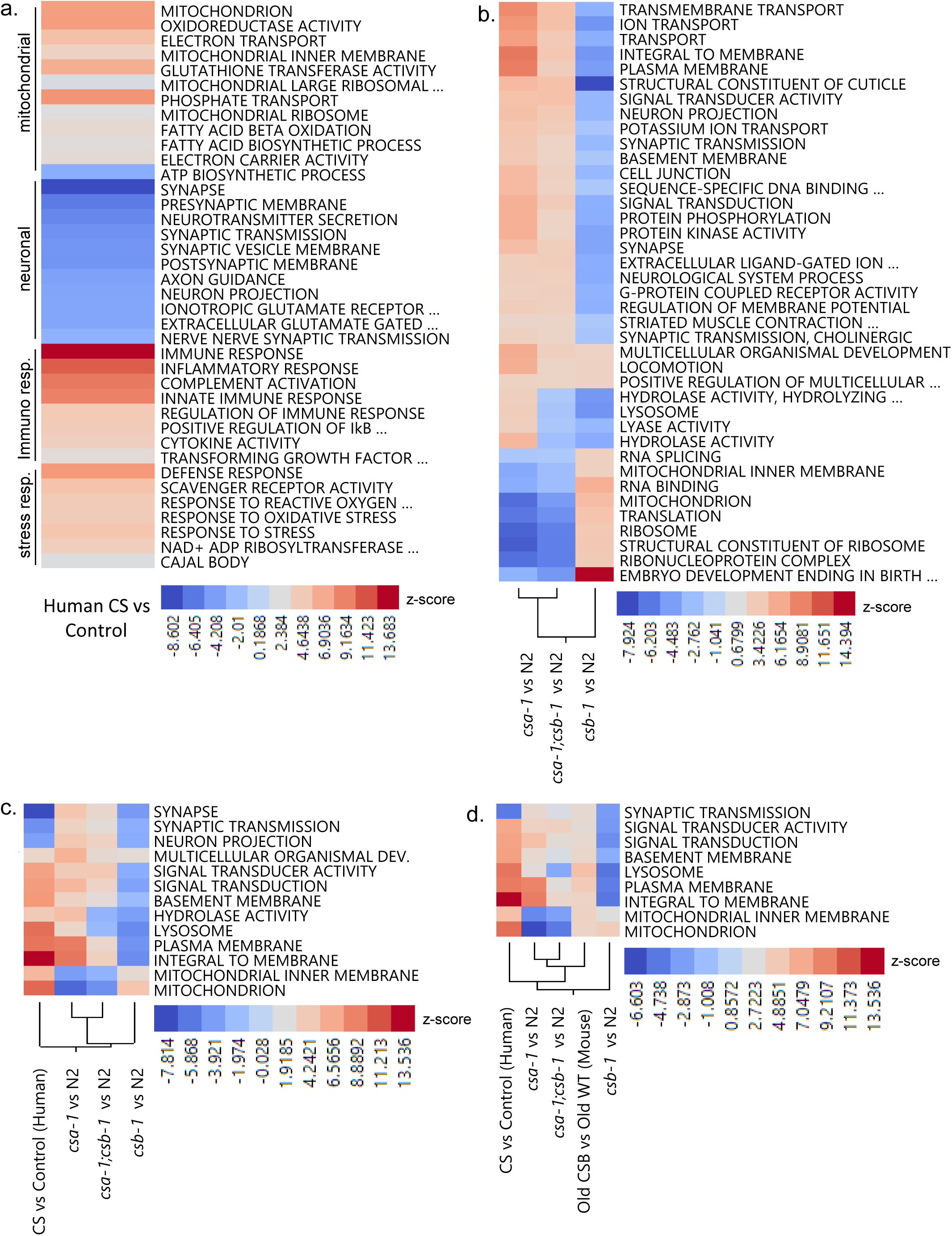
The transcriptomic analysis in CS postmortem brain tissue, CS mouse and *C. elegans* models. (**a**) Heatmaps of GO term Z-scores comparing human CS and control patients are shown. (**b**) Heatmaps of GO term Z-scores showing changes between CS worms compared to N2 worms. Hierarchical clustering showing (**c**) various GO terms commonly affected between human CS patient and CS worms and (**d**) GO terms that are commonly affected in CS mouse and worm models, and CS brain samples. A *p*-value ≤0.05 and an absolute value Z-Score cut-off of 2.0 were the cut-offs used for significance.

### Microarray analysis of Cockayne syndrome worm models

TC-NER is important for the DNA repair pathway in *C. elegans* [27], and mutations in both *csa-1* and *csb-1* lead to increased UV-sensitivity, suggesting that the function of these proteins is conserved in *C. elegans* [6, 28, 29]. We sought to evaluate *csa-1* and *csb-1* worm mutants to understand the shared or unique features of the disorder. Gene expression microarrays were performed on day 7 adult worms and compared to N2 worms to determine the set of significantly changed GO terms and how these compared to those observed in CS patients. After the selection of terms with *p*-values lower than 0.05 and an absolute value pathway Z-Score cut-off of 2.0, the remaining 39 GO terms and their respective Z-Scores were sorted by clustering in Fig 1b. Surprisingly, *csa-1* and *csb-1* worms had opposite pathway changes for a majority of the terms. Throughout the dataset, *csa-1;csb-1* worms partitioned more with *csa-1* than *csb-1* worms, suggesting that *csa-1* drives the double mutant phenotype. However, there was a shortlist of terms, including lysosome, where the *csa-1;csb-1* worms segregated with *csb-1* worms. Other terms in this group included histidine catabolic process to glutamate and formamide, oxidoreductase activity, and two terms related to hydrolase activity.

Clearly, the loss of CSB is dominant over CSA in these pathways. Previously, we reported that CS mice have an accumulation of dysfunctional mitochondria, with ribosomal and mitophagy defects [19]. These types of GO terms were significantly changed in the nematodes with mitochondrial and ribosomal pathways upregulated in *csa-1* and *csa-1;csb-1*, but down-regulated in *csb-1*. The lysosomal pathway, which is important for mitophagy [30], showed an opposite trend with up-regulation in *csa-1* but down-regulation in *csb-1* and *csa-1;csb-1* worms. The presence of changes in these GO terms, despite their direction, suggest the dysregulation in mitochondrial and ribosomal pathways in CS worms.

To understand the similar pathology in all CS models, we compared the GO terms in human CS patients (hCS) and CS worms. Only 13 GO terms, which were associated with mitochondria, synapse, membrane, and signal transduction, were in common (Fig. 1c). Interestingly, hierarchical clustering showed that *csb-1* worms cluster more closely to hCS than *csa-1* or *csa-1*;*csb-1* worms when the largest group of hCS terms listed in Figure 1a were compared (Supplementary Fig. 2). To understand the similar pathology in all CS models, we also included GO terms from the cerebellum of CSB mutant mice and their controls from a previously published study [4]. CSB mice recapitulate the features of small body size, hearing loss, retinal degeneration [31-33]. In addition to these features, we showed the presence of reduced muscle strength in old CSB mutant mice (Supplementary Fig. 3). We found that the terms associated with synaptic transmission, cellular signaling, membrane formation, lysosome, and mitochondria were the only GO terms that are commonly affected in all CS models and CS brain samples (Fig. 1d).

**Figure 2.**
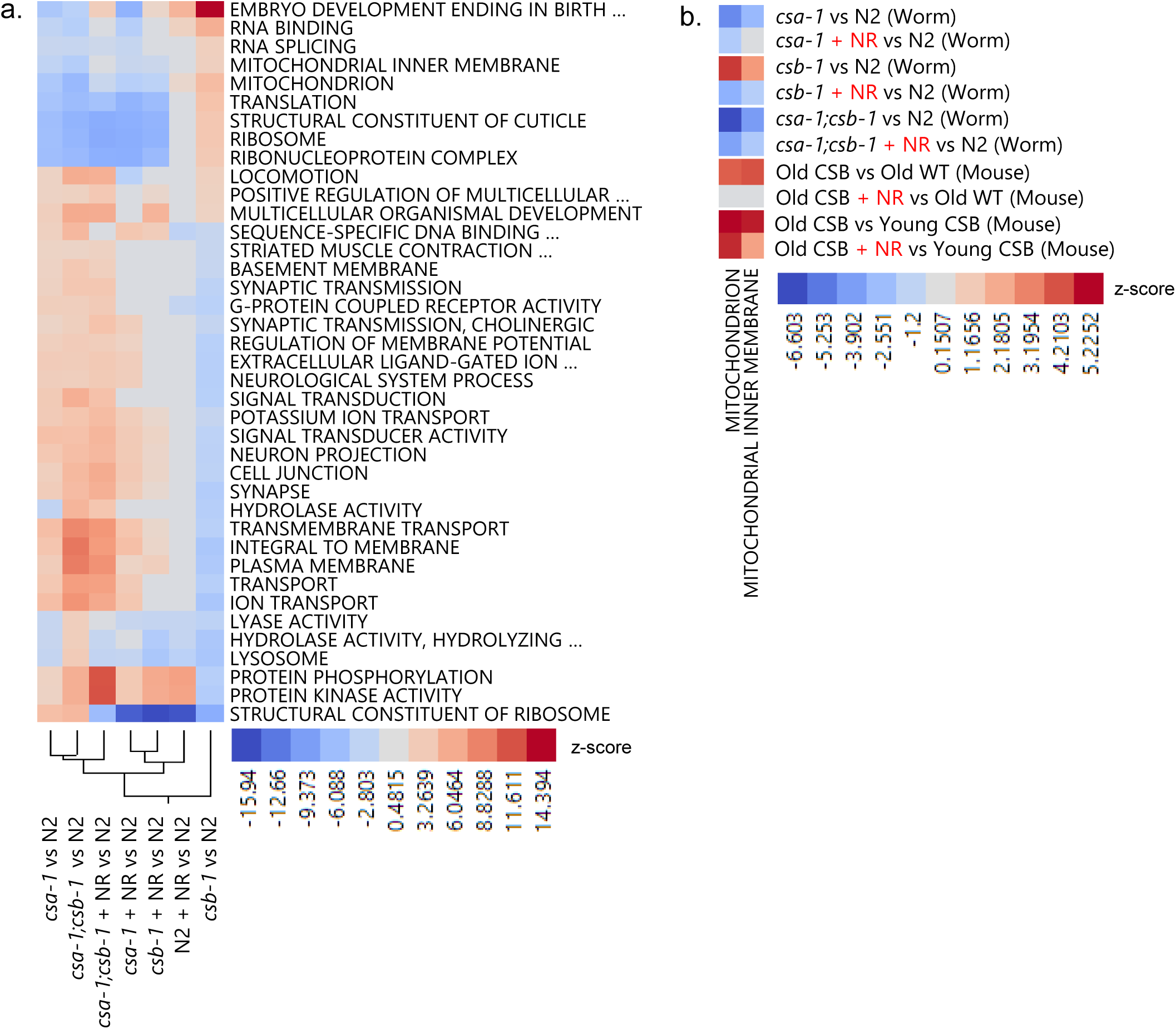
Transcriptomic analyses from CS *C. elegans* models and CS mouse without/with NR treatment. (**a**) Heatmap showing Z-score changes of the various GO terms from CS worms ± NR treatment (**b**) Heatmap showing Z-score changes of the GO terms related to mitochondria and mitochondrial inner membrane from CS worm and mouse models ±NR treatment. Terms were considered significant if they had *p*-values lower than 0.05 and an absolute value pathway Z-Score 2.0.

**Figure 3.**
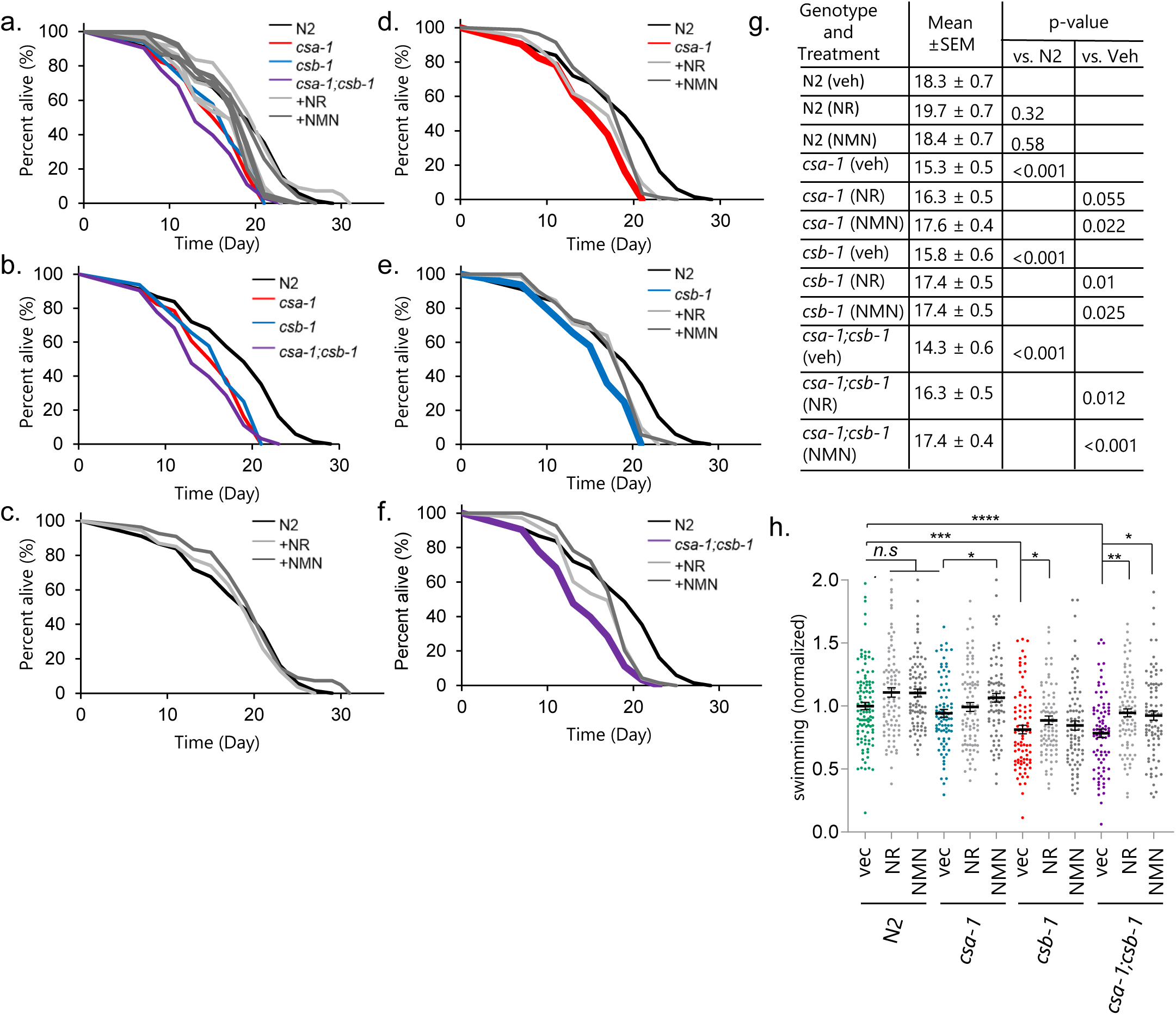
NAD^+^ replenishment rescues lifespan and healthspan of CS mutant *C. elegans*. (**a-f**) Effects of NAD^+^ supplementation on the lifespan of N2, *csa-1, csb-1*, and *csa-1*; *csb-1 worms* at 25□°C without/with 1□mM NR and NMN treatment. A representative set of data from two biological repeats is shown. The log-rank test was used for statistics, with *p* ≤ 0.05 was considered significant. (**g**) Tabular representation of lifespan data. (**h**) Effect of NR/NMN on swimming in N2 and CS worms at D8. Data are represented as mean□± □S.E.M (Total of 80-100 worms for each condition combined from three biologically independent experiments; Two-way ANOVA with Tukey’s post hoc test was used to determine significant differences. *p ≤ 0.05, **p ≤ 0.01, ***p ≤ 0.001, ****p ≤ 0.0001 and n.s., not significant.).

We previously showed that NAD^+^ supplementation with NR was of benefit in DNA repair-deficient mouse and worm models [16, 17]. Thus, we sought to determine if NR could modulate GO terms in this system as well. Microarray analysis was done on adult day 7 worms that were treated with vehicle (veh) or NR beginning the late L4 stage. Each genotype was compared to N2 vehicle worms for consistency. Overall, NR treatment modulated the gene expression profile in CS and caused CS mutant nematodes to cluster more closely with N2. To assess the pathways that are specially targeted by NAD^+^ supplementation, we examined the Z-score of individual GO terms that were altered in CS models but restored with NR treatment. The terms mitochondrion and mitochondrial inner membrane were the only common GO terms in CS mice and nematodes with Z-Score values that were normalized after NR treatment. Altogether, our comprehensive transcriptomic analysis suggests that the alterations in mitochondria play a central role in CS pathology and are conserved across animal models of CS.

### NAD^+^ replenishment in CS nematodes

The role of NAD+ supplementation in improving mitochondrial health was previously reported [16, 17, 34, 35]. We thus wondered whether targeting the pathological mitochondrial phenotype via NAD^+^ supplementation can alleviate disease phenotypes.

To address this, *csa-1, csb-1*, and *csa-1;csb-1* worms were analyzed for mean and maximum lifespan following NAD^+^ supplementation using two NAD^+^ precursors, NR (1mM) or NMN (1mM), from L4 stage (Fig. 3a-g). In the absence of drug treatment, N2 worms lived for 18.3 ±0.7 days while all the mutants showed reduced lifespans. *csa-1* showed 16% reduction (15.3 ±0.5 days), *csb-1* 13.6% reduction (15.8 ±0.6 days), and *csa-1;csb-1* 22% reduction (14.3 ±0.06 days). After drug treatments, in comparison to vehicle controls, both NAD^+^ supplement-treatment groups showed significantly improved lifespan of each genotype (Fig. 3g). No significant benefit of either drug was observed in N2 worms.

We next assessed healthspan and found that swimming motions were reduced all CS genotypes except the *csa-1* strain which displayed a slight reduction in swimming, but it did not reach significance (Fig. 3h). Notably, NR or NMN treatment significantly improved swimming at day (D) 8 in all CS genotypes (Fig. 3h). The pumping rate or maximum velocity was similar across all genotypes at the ages of 6 and 9 days (Supplementary Fig. 5). Collectively, all the worm data suggest that NAD^+^ augmentation extends lifespan and improves some of the healthspan parameters in the *csb-1*, and *csa-1;csb-1* worms.

### Mitochondrial assessment of CS nematodes

The GO term mitochondrion was one of the most significantly changed terms when comparing CS to N2 worms (Fig. 1-2). Along with the mitochondrial changes, the GO term for the lysosome was significantly changed. Mitophagy is a regulated process that involves the degradation of damaged mitochondria by lysosomes. This process plays an important role in mitochondrial homeostasis, neuroprotection, and even healthy longevity in laboratory animal models [36, 37]. While we have characterized mitophagy in the CSB^m/m^ mice [19], the changes of mitochondrial morphology and mitophagy in the animal models with CSA mutation or CSA/CSB double mutations are not known. We next sought to characterize mitochondrial networking and mitophagy in the CS *C. elegans* strains with or without NR treatment. In order to visualize the mitochondrial networking in the *C. elegans* body wall muscle tissue, we crossed *csa-1, csb-1*, and *csa-1;csb-1* mutants to a *myo3::gfp* reporter strain (N2 background) in which GFP is targeted to the mitochondria and the nuclei of body wall muscle cells. For these experiments, late L4 stage worms were treated with 1 mM NR. At day 7, worms were immobilized using Levamisole (100 mM) and imaged under a confocal microscope. All CS worm mutants exhibited significantly diminished levels of mitochondrial networking relative to N2 veh (Fig. 4a-b) and this phenotype was rescued with NR treatment in *csa-1* and *csb-1* strains. To directly visualize mitophagy in worms, the strains were crossed with DCT-1::GFP and LGG-1::RFP expressing worms [36]. Colocalization of the two markers is indicative of mitophagy and from this we can derive a ‘mitophagy score’. On day 6, *csb-1* and *csa-1;csb-1* worms displayed reduced mitophagy compared to N2 worms, by 28 and 23 percent, respectively, while there were no significant differences on day 1 (Fig. 4c and Supplementary Fig. 6). Notably, all CS genotypes exhibited a progressive reduction of mitophagy upon aging compared to day 1 while there was only a trend towards a reduction in the N2 worms (Fig. 4c, compare mitophagy score values on Day 1 to Day 6). To gain more insight into mitochondrial health in CS, we next assessed mitochondrial morphology in the CS strains by electron microscopy (Fig. 4d-e). Consistent with defective mitophagy in the CS worms, we observed increased mitochondrial length in all CS genotypes, which was restored with NR treatment. Notably, we found that NAD^+^ supplementation also restored increased mitochondrial width in *csb-1 and csa-1;csb-1* strains and mitochondrial area in *csa-1 and csa-1;csb-1* strains.

**Figure 4.**
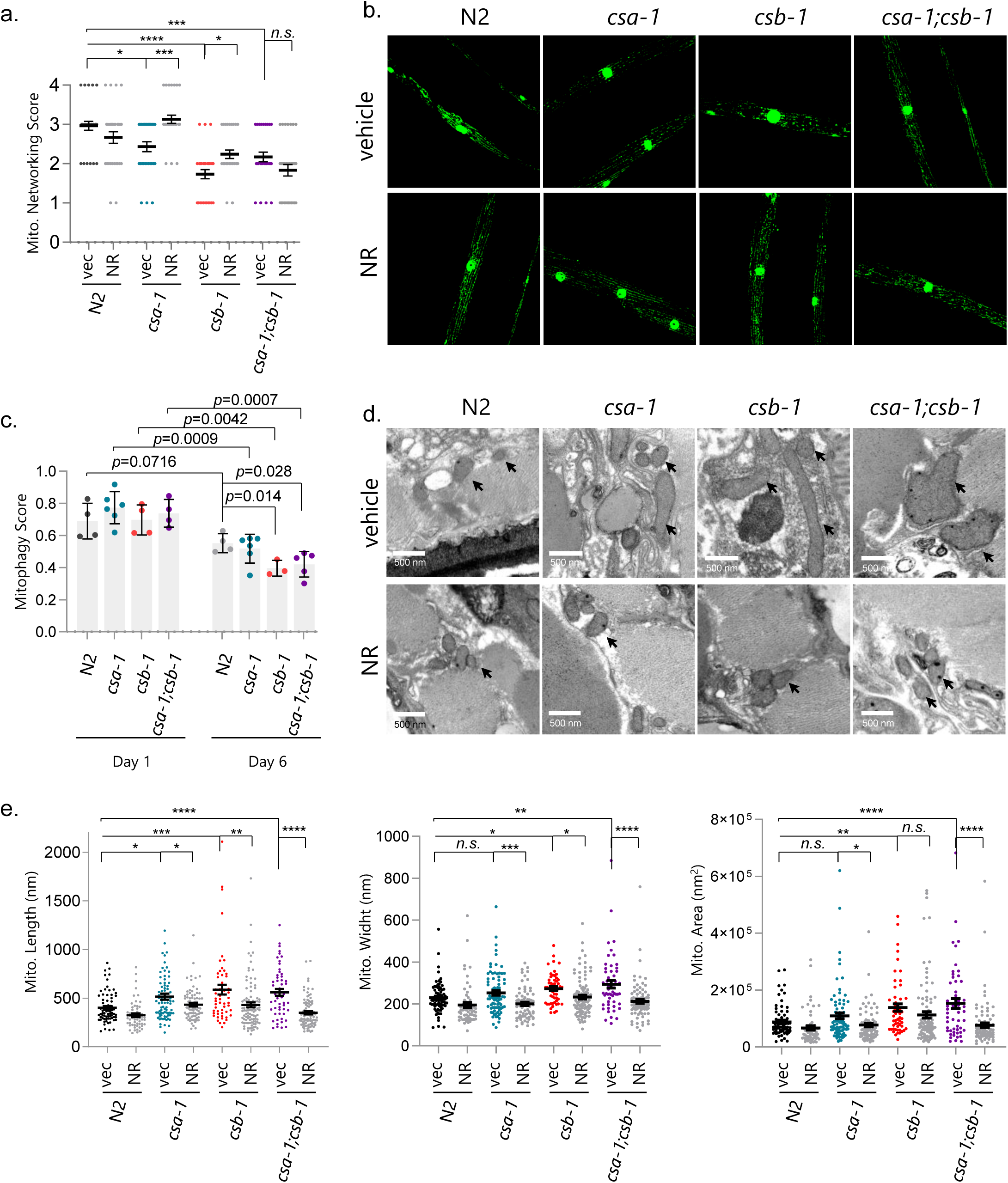
CS worms have reduced mitochondrial networking and mitophagy. (**a**,**b**) The *myo-3::gfp* reporter was expressed in both the nucleus and mitochondria to mark non-pharyngeal body wall muscle cells in the worms. Representative images and quantified scores of muscle mitochondrial morphology of adult D7 N2 and CS worms. Data are shown as mean□± □S.E.M (Total of 29-30 images of muscle cells from each group of worms (N=10), Two-way ANOVA with Tukey’s post hoc test was used to determine significant differences. *p ≤ 0.05, **p ≤ 0.01, ***p ≤ 0.001, ****p ≤ 0.0001 and n.s., not significant.). (**c**) The mitophagy reporter strain *N2;Ex(pmyo-3*∷*dsred*∷*lgg-1;pdct-1*∷*dct-1*∷*gfp)* was crossed with *csa-1, csb-1* or *csa-1*; *csb-1 worms* and imaged for muscle cells at D1 and D6 to measure the colocalization coefficient between DSRED∷DCT1 and LGG1∷GFP. Data are shown in mean□± □S.E.M (n□=□3-6 worms; Two-tailed *t*-test was used to determine significant differences.). (**d**) Mitochondrial structure changes were evaluated via electron microscopy (arrows indicate mitochondria). (**e**) Mitochondrial length, width, and area were quantified. Data are shown in mean□± □S.E.M (n□=□16-19 worms for each condition; Two-way ANOVA with Tukey’s post hoc test was used to determine significant differences. *p ≤ 0.05, **p ≤ 0.01, ***p ≤ 0.001, ****p ≤ 0.0001 and n.s., not significant.).

### Mitochondrial dynamics in CS patient cerebellum samples

The presence of impaired mitochondrial networking and morphology in CS worms (Fig. 4) suggests a dysregulation in mitochondrial homeostasis. We, therefore, assessed CS patient cerebellum samples for the activity of the proteins that have a key role in the turnover or dynamics of mitochondria (e.g. AMPK, DRP-1, and ULK1). AMPK directly phosphorylates downstream targets like PGC1α, which is important for mitochondrial biogenesis, and ULK1 (autophagy protein unc-51 like autophagy activating kinase 1) which is critical for mitophagy and thereby promotes mitochondrial homeostasis. We consistently observed a reduction in the activation of AMPK and its downstream target, ULK1, in CS patient samples (Fig. 5a and 5b). Further, the levels of Beclin1 were also decreased, suggesting a defect in the initiation of cellular autophagy in CS. In line with the findings of enlarged mitochondria in CS nematodes, we observed a trend towards a reduction in pDRP1 activity, a key protein of mitochondrial fission, in CS brain samples (Fig. 5). As previously reported [38], the levels of Gamma-H2AX, a marker of DNA damage, were increased in CS. The response to oxidative stress was upregulated in the microarray so we also tested for the reactive oxygen detoxifying enzymes Superoxide dismutase 1 (SOD1, cytoplasmic) and 2 (SOD2, mitochondrial). Although there was no significant change in SOD1 levels, we found that SOD2 levels increased in CS.

**Figure 5.**
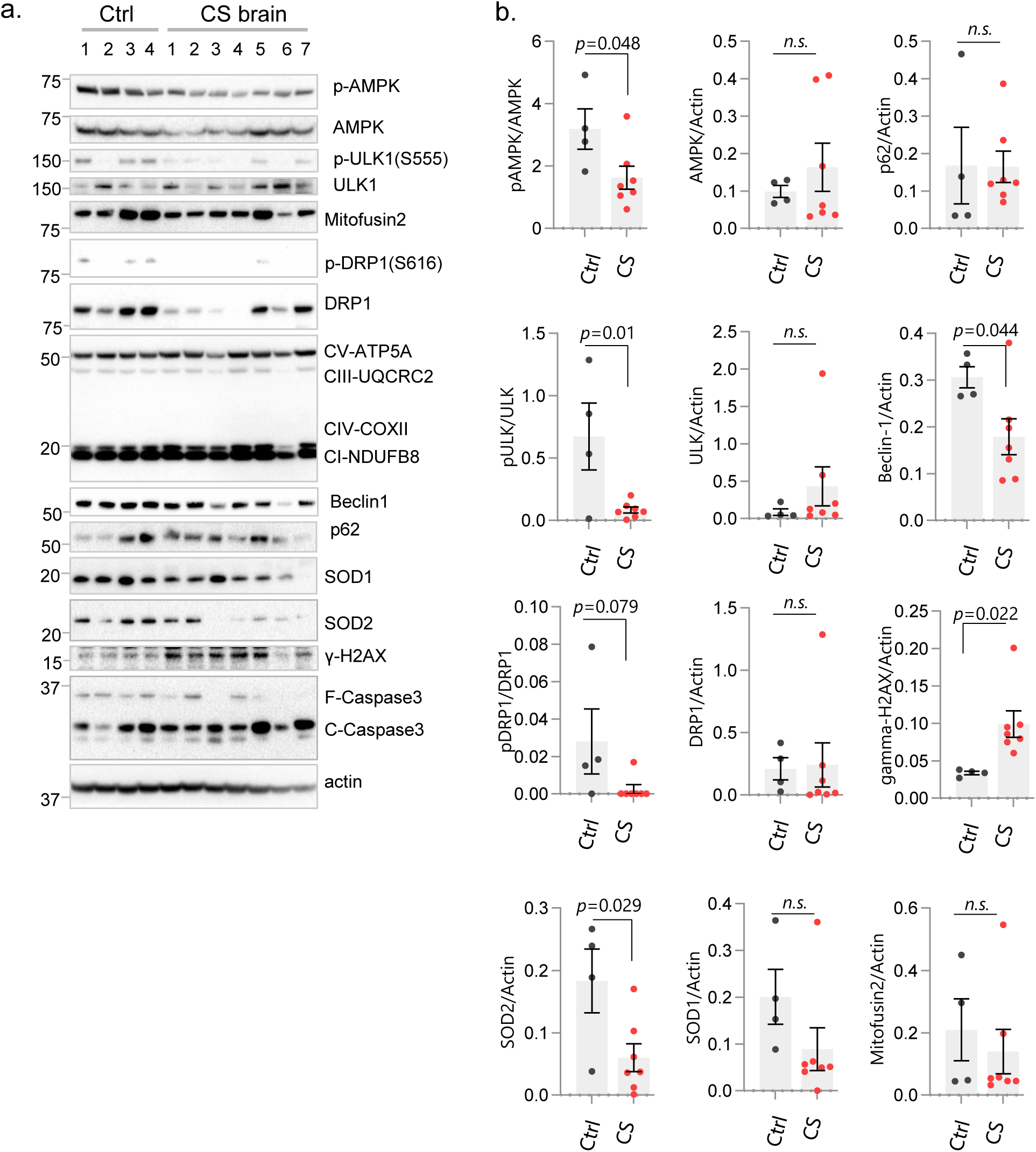
Protein expression levels in CS patient cerebellum samples. (**a**) Western blot data showing changes in the expression of designated proteins from control and CS patient cerebellum samples. The characteristics of the patient material are shown in Supplementary Table 3. (**b**) Graphical representation of statistical analysis of western blot data in *(a)* Data are shown in mean□± □S.E.M (n□=□3-6 worms; Two-tailed *t*-test was used to determine significant differences.).

### Restoration of AMPK signaling in CSA- and CSB-depleted fibroblasts with NAD^+^ supplementation

The effect of AMPK on NAD^+^ metabolism was previously published [39]. To investigate whether NAD^+^ supplementation could modulate the expression and activation of mitophagy and autophagy-related proteins, we treated primary fibroblasts with NMN following CSA or CSB depletion. In line with the findings in CS brain samples, we observed lower levels of activated AMPK (p-AMPK), pULK1 and pDRP1 in CSA and CSB deficient primary fibroblasts (Fig. 6). Notably, NAD^+^ supplementation restored phosphorylation of these proteins, which is predicted to increase their activity (Fig. 6). Beclin1 levels were slightly lower in CSA and CSB depleted cells compared to control cells but increased significantly following NMN treatment. The p62 protein levels in CSA and CSB deficient cells show no difference from the control group but decrease following NMN treatment in all samples, suggesting that cellular autophagy is increased after NAD^+^ supplementation. Collectively, these findings suggest that impaired activation of AMPK and ULK1 along with DRP1 leads to dysregulation of mitochondrial fission and degradation and causes subsequent defects in mitochondrial homeostasis. NAD^+^ supplementation restores these features and improves mitochondrial quality in CS.

**Figure 6.**
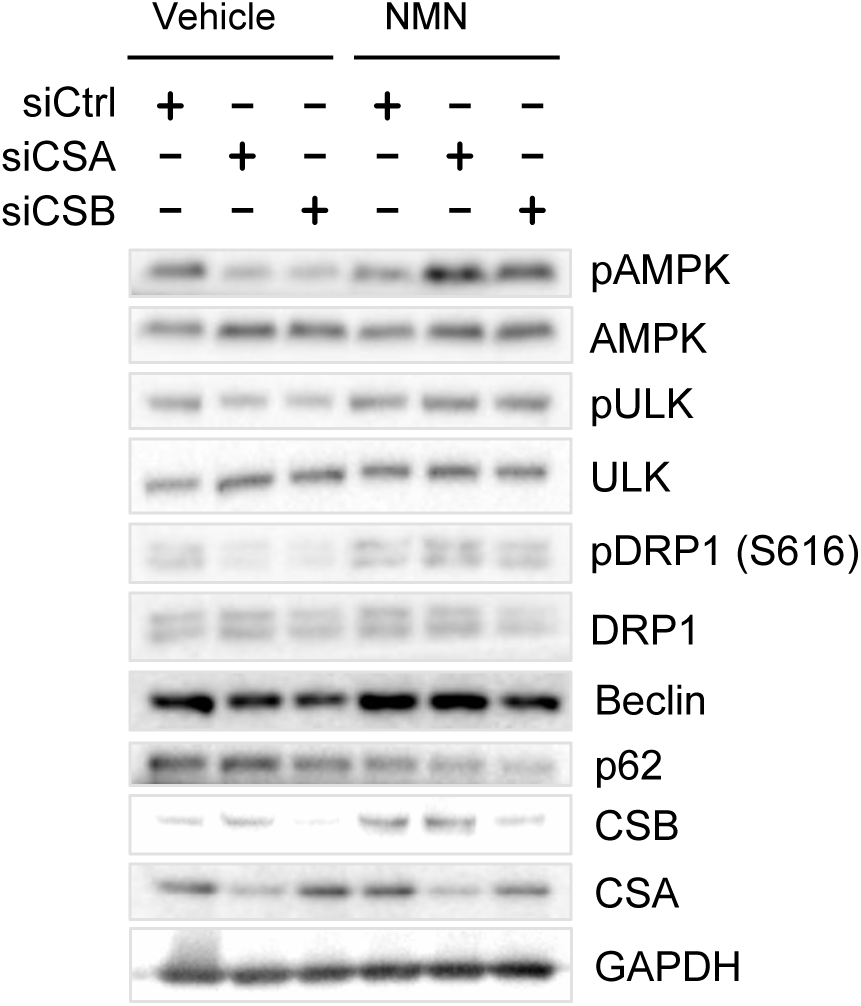
NAD^+^ supplementation rescues AMPK signaling in CSA and CSB depleted fibroblasts. Western blot data showing changes in the expression of designated proteins from control and CSA- or CSB-depleted primary fibroblast cell line (passage 9).

## Discussion

Mitochondrial homeostasis involves coordination between mitochondrial biogenesis and mitophagy. If these processes become uncoordinated, mitochondrial dysregulation can ensue. Hence, mitochondrial dysfunction is causative in many human ailments, including neurological abnormalities [40, 41]. It is important to investigate common phenotypes between human and animal models of CS to explore conserved pathways. Here, we report that mitochondrial dysfunction is a common feature across CS human and animal models. Comparative transcriptomic analysis of CS patient cerebellum samples and CS animal models like *C. elegans* and mice converge to implicate mitochondrial dysfunction in CS. NAD^+^ depletion is related to multiple hallmarks of aging [42] and NAD^+^ repletion ameliorates some of these phenotypes in CS mouse models [4]. Here, we observed that NAD^+^ supplementation improves healthspan and lifespan in CS worm models as well. Also, we showed that NAD^+^ supplementation enhances mitochondrial networking, stimulates mitophagy and leads to a normalization of mitochondrial morphology via restoration of increased length. Specifically, mitochondrion and mitochondrial inner membrane GO terms were the only GO terms that NR treatment nearly reverted to wild type in both CS mouse and worm models, suggesting NR’s therapeutic effects are mediated through mitochondrial homeostasis in CS.

Our data suggest that impairment of neuronal pathways along with mitochondrial dysregulation contributes to the neurodegeneration in CS. CSB has been known to localize to mitochondria and participate in DNA repair and transcription [8, 20, 21, 43]. Neurological degeneration is observed in many conditions with DNA repair deficiency including CS [1]. Cells with compromised mitophagy contribute to neuronal death and thus cause neurodegeneration [44]. Our recent report suggests that NAD^+^-dependent SIRT1 activity regulates mitophagy through a DAF-16-DCT1 pathway [17]. AMPK is associated with NAD^+^ metabolism and SIRT1 activity, and its activation regulates ULK1/DRP1 proteins necessary for mitophagy [45, 46]. Protein expression from the brain samples of CS patients showed dysregulation of AMPK and decreased phosphorylation of its downstream target, ULK1, which is necessary for ULK1’s translocation to damaged mitochondria for mitophagy. Although we did not observe a significant reduction in mitofusin2, there was a trend toward reduction in phosphorylated dynamin-like protein DRP1, which mediates the mitochondrial fission and subsequent fragmentation to undergo mitophagy [47]. This finding is in line with the increased mitochondrial length in all CS worm strains, suggesting defects in mitochondrial fragmentation. AMPK helps in recruiting DRP1 to the mitochondrial outer membrane and thus regulates the morphology of mitochondria [46, 48]. This imbalanced regulation of mitophagy was further supported by the increased levels of DNA damage and ROS [49]. Interestingly, we noted that NMN treatment helps regulate AMPK pathway proteins. Thus, in CS, increased DNA damage causes increased PARP-1 mediated NAD^+^ consumption, thereby leading to depletion in the NAD^+^ pool [4]. This NAD^+^ pool is regulated by the AMPK-ULK1/DRP1 pathway whereas dysregulation of CS proteins generates a mitochondrial dysfunction phenotype in CS (Fig. 7). The extracellular NAD^+^ degradation produces increased adenylate pool in cells and increases the intracellular ATP levels by activating AMPK through modulation of the ATP/AMP ratio [50]. Thus, AMPK senses stress to initiate mitochondrial fission to eliminate damaged mitochondria while at the same time induce mitochondrial biogenesis to regulate mitochondrial homeostasis [48]. Given the central role of NAD^+^ metabolism and mitochondria in energy production, CSA and CSB play an important role to modulate the AMPK-ULK1/DRP1 pathway to maintain mitochondrial homeostasis.

**Figure 7.**
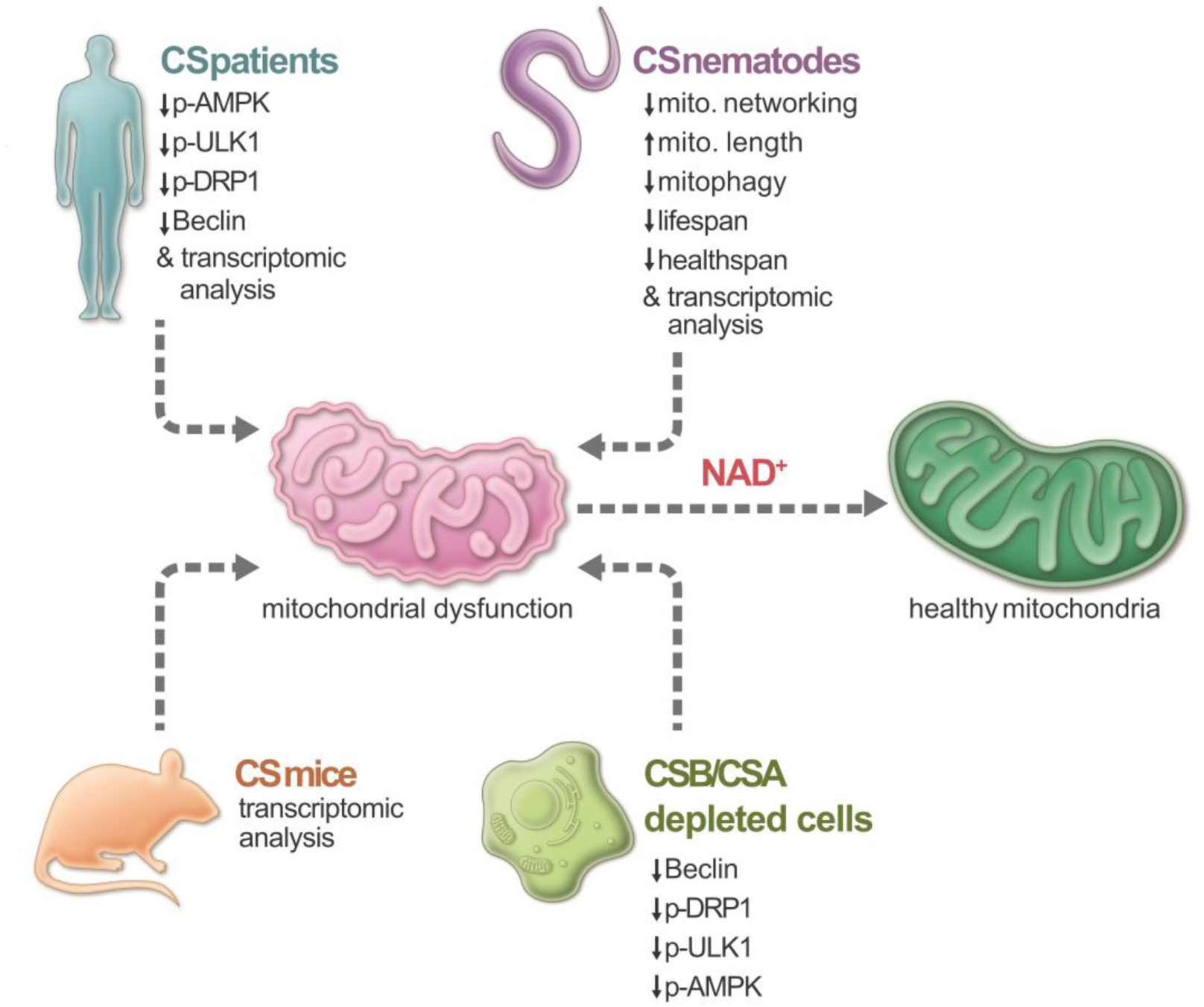
We propose CS mutation leads to cellular NAD^+^ reduction that impairs mitochondrial homeostasis through downregulation of the activities of p-AMPK and p-ULK1. NAD^+^ levels decline during the aging process, resulting in many age-associated pathologies like mitochondrial dysfunctions. Restoring NAD^+^ level by supplementing NAD^+^ precursors such as NR (nicotinamide riboside) and NMN (nicotinamide mononucleotide) can ameliorate these age-associated defects.

## Conclusions

In closing, these results illustrate the underlying molecular mechanisms of CS proteins (CSA and CSB) against premature age-induced mitochondrial dysfunction. Moreover, the interplay between NAD^+^ supplementation, mitochondria, and mitophagy provides insight into the development of therapeutic strategies in the treatment of DNA repair accelerated aging models like CS.

## Methods

### *C. elegans* strains and cultivation

All *C. elegans* strains were maintained and grown on standard nematode growth medium (NGM) as previously described [51]. Bristol strain N2 (wild type) and *myo3::gfp* reporter strains were obtained from the Caenorhabditis Genetics Center (CGC). Worm populations were synchronized via egg lay and all experiments were conducted at 25°C unless otherwise noted. *csa-1* (tm4539) and *csb-1* (ok2335) were provided by Dr. Bjorn Schumacher (University of Cologne, Cologne, Germany). The mitophagy reporter strain *N2;Ex(pmyo-3*∷*dsred*∷*lgg-1;pdct-1*∷*dct-1*∷*gfp)* was provided by Dr. Nektarios Tavernarakis (Institute of Molecular Biology and Biotechnology, Crete, Greece) and described previously [36].

### *C. elegans* drug treatment

Worms were treated with 100 µM Fluorodeoxyuridine (FUDR) in order to maintain only adult worms throughout the study (unless otherwise noted), as described previously [52]. Worms were transferred manually to plates containing FUDR at the late L4 developmental stage. Worms were treated with NAD^+^ precursors nicotinamide riboside (NR) and nicotinamide mononucleotide (NMN) at a final concentration of 1 mM in water. Drug plates changed every three days. Water was used as vehicle control. The drug was added directly to seeded plates one day ahead of the experiments. Worms were then transferred to the treatment plates by hand.

### *C. elegans* lifespan analysis

Lifespan analysis was conducted as described previously [16]. Worm populations were synchronized via egg lay and transferred to treatment plates at the L4 stage (48 hours post egg lay at 25°C). Experiments were conducted at 25°C unless otherwise noted. Animals were scored every 2 days. Worms that were nonresponsive to touch were considered dead. Missing worms or worms that died due to matricidal hatching, desiccation, or gonad extrusion were censored. Statistical significance was determined using the log-rank test. *p* ≤ 0.05 was considered significant. The data is represented by a smoothed Kaplan-Meier survival curve.

### *C. elegans* mitochondrial networking

To visualize the mitochondrial networking in the *C. elegans* body wall muscle tissue, *csa-1, csb-1*, and *csa-1;csb-1* mutants were crossed to a *myo3::gfp* reporter strain (N2 background). Images were acquired with a Zeiss confocal microscopy and analyzed as described previously [53]. Worms were transferred to drug-treated plates from the late L4 stages until Day 7 unless otherwise noted. Images were analyzed for mitochondrial networking integrity in a double-blinded manner. Mitochondrial networking was assessed using an arbitrary scale from 1-5. Healthy, organized mitochondrial networks in which the mitochondria are longer, less circular, aligned with the muscle fibers, and with reduced interconnecting mitochondrial strings received a score of 5 [53]. The mitochondrial networking score was reduced if the mitochondria appeared disorganized and more circular with mitochondria forming a network of irregular shapes. A total of 29-30 images of muscle cells from each group of worms (N=10) were collected.

### *C. elegans* healthspan analysis

Methods to quantify healthspan in *C. elegans* have been described previously [54-56]. Healthspan was assessed by maximum velocity, pharyngeal pumping, and swimming. To measure maximum velocity, we adapted a method previously described by Hahm et al [56]. On the designated day, up to 10 worms were transferred to unseeded NGM plates. The movement was recorded over a span of 30 seconds using a stereomicroscope (Unitron 2850) and a charge-coupled device camera (Infinity 2, Lumenera). Videos were analyzed using ImageJ software and the wMTrck Batch (http://www.phage.dk/plugins) plugin. To assess pharyngeal pumping, worms were assayed at the adult D6 stage. Contractions of the pharynx were manually counted over a period of 30s seconds in a double-blinded manner. At least 30 worms were measured/group and the experiments were conducted three separate times. Swimming analysis was conducted as described previously [53]. At D8, individual worms were transferred from treatment plates onto a 6 cm dish containing 1 mL of M9 buffer. After an acclimatization period of ∼10 s, the number of body bends was scored over a period of 30 seconds.

### *C. elegans* mitophagy

To visualize mitophagy in worms, the mitophagy reporter strain *N2;Ex(pmyo-3*∷*dsred*∷*lgg-1;pdct-1*∷*dct-1*∷*gfp)* was crossed with *csa-1, csb-1* and *csa-1*; *csb-1 worms*. The muscle cells were imaged at D1 and D6 under confocal microscopy as described previously [36]. A total of 3∼6 worms/group were imaged. Zeiss LSM Image Examiner was used to measure the colocalization coefficient between DSRED∷DCT1 and LGG1∷GFP.

### Electron microscopy (EM) for mitochondrial morphology

For electron microscopy (EM) of mitochondrial morphology of *C. elegans*, the worms were freshly dissected and fixed in 2% paraformaldehyde, followed by EM as described elsewhere [17].

### Cockayne syndrome cerebellum samples

Human tissues were obtained from the NIH NeuorBioBank’s Brain and Tissue repository at the University of Maryland, Baltimore. The patients and their ID numbers are listed in Supplementary Tables 1 and 3.

### Microarray analysis

RNA was purified from cerebellum samples from Cockayne patients and age- and gender-matched healthy individuals by using the Nucleospin RNA isolation kit (catalog no. 740955.250; Macherey-Nagel) with initial quantitation conducted using a NanoDrop ND-1000 spectrophotometer (Thermo Fisher Scientific). A 2100 Bioanalyzer (Agilent Technologies) was used to confirm the quality of the RNA. The microarray was performed by the Gene Expression and Genomics Unit core facility (NIA) and analyzed using DIANE 6.0 software. The complete set was tested for Geneset enrichment using parametric analysis of gene set enrichment (PAGE). Detailed data analysis was performed as reported previously [4].

For microarray analysis of nematodes, N2, *csa-1, csb-1, csa-1;csb-1* worms were exposed to 500 µM NR from the L4 stage and whole-body tissue was collected at adult day 7 for RNA extraction (Qiagen RNeasy mini kit with a QIAcube machine) [57]. Three biological replicates were prepared for each group. The Agilent kits were used for *C. elegans* microarray with the reagents including *C. elegans* (V2) Gene Expression Microarray (# G2519F-020186), Low Input Quick Amp Labeling Kit (one color, #5190-2305), Hybridization Backing Kit (# G2534-60012), and Gene Expression Hybridization Kit (#5188-5242). The microarray was performed and analyzed at the National Institute on Aging (NIA) Gene Expression and Genomics Unit using DIANE 6.0 software. Raw microarray data were log-transformed to yield z-scores. The z-ratio was calculated as the difference between the observed gene z-scores for the experimental and the control comparisons, divided by the standard deviation associated with the distribution of these differences. Genes whose Z-ratio was ± 1.5 with a p-value ≤0.05, 0.3 false discovery rate (FDR), and average signal intensity of comparison greater than zero. The gene expression z-ratio values were then used as input to perform Parametric Analysis of Gene Set Enrichment (PAGE) testing [58]. The accession number for both array data is GSE144558.

### Cell culture and cell lines

The normal human fetal lung primary fibroblasts were received from Coriell Institute (ID #I90-83) and were cultured in Dulbecco’s modified Eagle medium (DMEM) supplemented with 10% (vol/vol) FBS and 1% penicillin/streptomycin in a humidified chamber under 5% (vol/vol) CO2 at 37 °C.

### siRNA knockdown

siERCC6 (5′-CCACUACAAUAGCUUCAAGACAGCC-3′), siERCC8 (5′-GGAGAACAGAUAACUAUGCUUAAGG −3′) and siRNA duplex control (Origene) was diluted with DMEM to a final concentration of 20□nM, mixed with INTERFERin (Polyplus transfection), incubated for 15 min at room temperature and transfected into primary cells target according to the manufacturer’s instructions. 24 hours after transfection, the media was replaced with FBS media. 48 hours after transfection, the cells were treated with NMN (500uM) or water as a vehicle for 24 hours and lysed for western blotting.

### Immunoblotting

Cells were lysed in RIPA buffer (#9806S; Cell Signaling) with Halt™ Protease and Phosphatase Inhibitor Cocktail (100X) (#78444, ThermoFisher Scientific) unless indicated otherwise. Antibodies were used according to the manufacturer’s instructions to detect the following antigens: AMPK (Cell signaling, #5831), pAMPK (Thr172) (Cell signaling, #2535), pULK1 (Ser555, Cell signaling, #5869), ULK1 (Cell signaling, #6439), p62 (Cell signaling, #39749), β-actin (Santa Cruz, #sc-1616), DRP1 (D8H5, Cell signaling, #5391) DRP-1 (Cell signaling, #8570), p-DRP-1 (S616, Cell signaling, #3455), Beclin-1 (Cell signaling, #3738), mitofusin 2 (Cell signaling, #9482), total OXHOS rodent cocktail (Abcam, ab110413), SOD-1 (Cell signaling, #2770), SOD-2 (Enzo Life Sciences, #AD1-SOD-110-F), γ-H2AX (Santa Cruz, #sc-517348), GAPDH (Abclonal, #AC027).

### Animals

Mouse models of CSB^m/m^ (mice carrying a premature stop codon in exon 5 to mimic the K337 stop truncation mutation from CS patient (CS1AN) [7] and wild-type (WT) on a C57BL/6J background in the age ranges of 20-26 weeks (young), 52-62 weeks (mid-age), and 98-11 weeks (old) of age were used for the grip-strength studies. All animal protocols were approved by the appropriate institutional animal care and use committee of the National Institute on Aging.

### Grip Strength

The forelimbs of each mouse were tested for grip strength by pulling on a wire attached to a Chatillon DFE002 force gauge (Chatillon Force Measurement Systems). Five pulls were performed for each mouse and the mean of the recordings was determined as a final score.

### Statistical analysis

All statistical analyses were performed with GraphPad Prism (GraphPad Software, Inc.). For multiple samples with two groups, two-way ANOVA with Tukey’s post hoc test was used to determine significant differences. One-way ANOVA with Tukey’s post hoc test was used to determine significant differences across multiple samples. T-test was used to compare two groups. The log-rank test was used to determine statistical significance for survival assays and *p* ≤ 0.05 was considered significant.

## Declarations

### Ethics approval and consent to participate

All animal protocols were approved by the appropriate institutional animal care and use committee of the National Institute on Aging. Human tissues were obtained from the NIH NeuorBioBank’s Brain and Tissue repository at the University of Maryland, Baltimore.

## Consent for publication

Not applicable

## Availability of data and materials

The datasets used and/or analysed during the current study are available from the corresponding author on reasonable request.

## Authors’ contributions

E.F.F. and M.N.O. planned the experiments. E.M.F., M.N.O., E.F.F., and V.T. performed the experiments; V.A.B. and D.L.C supervised the findings of the work; D.L.C, M.N.O. and V.T. wrote the manuscript with support from E.M.F.; All authors discussed the results and contributed to the final manuscript.

## Acknowledgments

We would like to thank Drs. Kevin Becker, Yongqing Zhang and Elin Lehrmann, and the Gene Expression and Genomics Unit, Laboratory of Genetics and Genomics, NIA IRP for performing microarray analyses. Dr. Elin Lehrmann performed all validation, experiments, and data extraction for array experiments and the GEO submission while Dr. Yongqing Zhang performed the bioinformatic analysis. We would like to thank Ms. Wendy B. Iser, Mr. David M. Figueroa, Mr. Mark Wilson, Mr. Jesse Kerr, and Dr. Henok Kassahun for their help in performing worm-related experiments. We also thank Dr. Bjorn Schumacher (University of Cologne, Cologne, Germany) for providing *csa-1* (tm4539) and *csb-1* (ok2335) strains. We thank Drs. Yujun Hou and Louise Christiansen for their critical reading of the manuscript.

## Supplementary Table Legends

**Supplementary Table 1**. The list of Cockayne Syndrome patient cerebellum samples that were used for microarray analysis.

**Supplementary Table 2**. Top 100 upregulated and downregulated GO terms in Cockayne Syndrome patient cerebellum samples. An absolute value pathway Z-score cut-off of 2.0 and *p*-values ≤0.05 were considered significantly changed.

**Supplementary Table 3**. The list of Cockayne Syndrome patient cerebellum samples that were used for western blot analysis.

## Supplementary Figure Legends

**Supplementary Figure 1**. Clustering of patient samples from CS-derived patient cerebellum samples and control-derived samples.

**Supplementary Figure 2**. Clustering of GO terms from CS-derived patient cerebellum and worm samples. An absolute value pathway Z-score cut-off of 2.0 and *p*-values ≤0.05 were considered significantly changed.

**Supplementary Figure 3**. The grip strength of each mouse was scored as described in the methods section. Data are shown in mean□±□S.E.M (n□=□5 for each group; 2-way ANOVA with Fisher LSD test was used to determine significant differences). The age range is 20-26 weeks for young mice, 52-62 weeks for mid-age mice, and 98-11 weeks for old mice.

**Supplementary Figure 4**. Heatmap showing how GO terms related to mitochondria and mitochondrial inner membrane are changed ±NR from CS human, worms and mouse comparisons without/with NR treatment.

**Supplementary Figure 5**. The Effect of NR/NMN on pharyngeal pumping at D6 (**a**) and maximum velocity at D9 (**b**) in N2 and CS worms. Data are represented as mean□± □S.E.M. (Total of 175-195 worms for each condition combined from three biologically independent experiments for velocity assessment; a total of 40-140 worms for each condition combined from two biologically independent experiments for velocity assessment; Two-way ANOVA with Tukey’s post hoc test was used to determine significant differences.

**Supplementary Figure 6**. Representative images showing colocalization between DSRED∷DCT1 and LGG1∷GFP in muscle cells in worms.

