## supplemental data for "Cockayne syndrome proteins CSA and CSB maintain mitochondrial homeostasis through NAD^+^ signaling"

### Supplementary Figures and Table

### Supplementary Table 1

Cockayne Syndrome patient cerebellum samples that were used for microarray analysis.

| Group | UMBN | Age | Sex | Cause of Death | PMI |
|---|---|---|---|---|---|
| Control (WT1) | 3415 | 7 | M | Multisystem Failure | 12 |
| Control (WT3) | 5976 | 4 | F | Smoke Inhalation | 21 |
| Control (WT4) | 5928 | 27 | F | Cardiac arrhythmia | 26 |
| Control (WT5) | 1078 | 17 | F | Accident, Multiple Injuries | 12 |
| Control (WT6) | 5391 | 8 | M | Drowning | 12 |
| Cockayne (CS1) | 1920 | 7 | M | Diffused alveolar damage / acute pneumonia | 16 |
| Cockayne (CS2) | 786 | 17 | F | Complication of Disorder | Unknown |
| Cockayne (CS3) | 5492 | 7 | M | Complications of Disorder | Unknown |
| Cockayne (CS4R) | 5581 | 27 | F | Complication of Disorder | 14 |
| Cockayne (CS6) | 1124 | 4 | F | Complications of Disorder | 3 |
| Cockayne (CS7) | 5105 | 8 | M | Complication of Disorder | 6 |
| Cockayne (CS8) | 1762 | 4 | F | Complications of Disorder | 8 |

The list of Cockayne Syndrome patient cerebellum samples that were used for microarray analysis.

### Supplementary Table 2

#### Top 100 upregulated and downregulated GO terms in CS patient cerebellum samples

| Gene Ontology Term | Zscore |
|---|---|
| GO0005615 EXTRACELLULAR SPACE | 17.1982877 |
| GO0016020 MEMBRANE | 14.3827928 |
| GO0006955 IMMUNE RESPONSE | 13.89056137 |
| GO0016021 INTEGRAL TO MEMBRANE | 13.5362495 |
| GO0046870 CADMIUM ION BINDING | 11.90484511 |
| GO0007155 CELL ADHESION | 11.54209473 |
| GO0006954 INFLAMMATORY RESPONSE | 10.94812908 |
| GO0005737 CYTOPLASM | 10.63895048 |
| GO0005520 INSULIN LIKE GROWTH FACTOR BINDING | 10.42750915 |
| GO0005739 MITOCHONDRION | 10.18613769 |
| GO0005764 LYSOSOME | 9.85877125 |
| GO0006956 COMPLEMENT ACTIVATION | 9.840334507 |
| GO0045087 INNATE IMMUNE RESPONSE | 9.809865555 |
| GO0005576 EXTRACELLULAR REGION | 9.735861816 |
| GO0005886 PLASMA MEMBRANE | 9.581756988 |
| GO0005578 PROTEINACEOUS EXTRACELLULAR MATRIX | 9.360849996 |
| GO0005178 INTEGRIN BINDING | 9.303586999 |
| GO0016491 OXIDOREDUCTASE ACTIVITY | 9.036500462 |
| GO0005887 INTEGRAL TO PLASMA MEMBRANE | 8.849482637 |
| GO0005515 PROTEIN BINDING | 8.809728894 |
| GO0019838 GROWTH FACTOR BINDING | 8.597717018 |
| GO0006952 DEFENSE RESPONSE | 8.500907053 |
| GO0005783 ENDOPLASMIC RETICULUM | 8.394284999 |
| GO0004872 RECEPTOR ACTIVITY | 8.326364494 |
| GO0005509 CALCIUM ION BINDING | 8.298073392 |
| GO0006817 PHOSPHATE TRANSPORT | 8.237169775 |
| GO0006935 CHEMOTAXIS | 8.127819547 |
| GO0008152 METABOLIC PROCESS | 8.083951694 |
| GO0005581 COLLAGEN | 7.883409182 |
| GO0005507 COPPER ION BINDING | 7.826236993 |
| GO0005604 BASEMENT MEMBRANE | 7.745480057 |
| GO0007165 SIGNAL TRANSDUCTION | 7.556848936 |
| GO0005198 STRUCTURAL MOLECULE ACTIVITY | 7.537305994 |
| GO0030020 EXTRACELLULAR MATRIX STRUCTURAL CONSTITU | 7.308711423 |
| GO0004871 SIGNAL TRANSDUCER ACTIVITY | 7.174696738 |
| GO0001525 ANGIOGENESIS | 7.174292709 |
| GO0016337 CELL CELL ADHESION | 7.058800126 |
| GO0007160 CELL MATRIX ADHESION | 6.93721382 |
| GO0008284 POSITIVE REGULATION OF CELL PROLIFERATIO | 6.904929478 |
| GO0001726 RUFFLE | 6.709941735 |
| GO0006928 CELL MOTILITY | 6.697710283 |
| GO0007166 CELL SURFACE RECEPTOR LINKED SIGNAL TRAN | 6.61859733 |
| GO0003779 ACTIN BINDING | 6.522361001 |
| GO0001558 REGULATION OF CELL GROWTH | 6.518922008 |
| GO0004364 GLUTATHIONE TRANSFERASE ACTIVITY | 6.516375942 |
| GO0006915 APOPTOSIS | 6.500390316 |
| GO0001667 AMEBOIDAL CELL MIGRATION | 6.377888472 |
| GO0003924 GTPASE ACTIVITY | 6.246543791 |
| GO0009615 RESPONSE TO VIRUS | 6.235797044 |
| GO0006118 ELECTRON TRANSPORT | 6.205399656 |
| GO0005201 EXTRACELLULAR MATRIX STRUCTURAL CONSTITU | 6.170621769 |
| GO0008283 CELL PROLIFERATION | 6.157249716 |
| GO0048514 BLOOD VESSEL MORPHOGENESIS | 6.036897874 |
| GO0003824 CATALYTIC ACTIVITY | 6.005319953 |
| GO0006629 LIPID METABOLIC PROCESS | 6.002842716 |

| Gene Ontology Term | Zscore |
|---|---|
| GO0004866 ENDOPEPTIDASE INHIBITOR ACTIVITY | 5.995035231 |
| GO0006916 ANTI APOPTOSIS | 5.933099193 |
| GO0004180 CARBOXYPEPTIDASE ACTIVITY | 5.87220924 |
| GO0005856 CYTOSKELETON | 5.861923353 |
| GO0005829 CYTOSOL | 5.803244589 |
| GO0030593 NEUTROPHIL CHEMOTAXIS | 5.75391386 |
| GO0015629 ACTIN CYTOSKELETON | 5.709100582 |
| GO0008360 REGULATION OF CELL SHAPE | 5.706752579 |
| GO0042632 CHOLESTEROL HOMEOSTASIS | 5.704322613 |
| GO0005743 MITOCHONDRIAL INNER MEMBRANE | 5.69476918 |
| GO0005625 SOLUBLE FRACTION | 5.674069526 |
| GO0030036 ACTIN CYTOSKELETON ORGANIZATION AND BIOG | 5.628093311 |
| GO0001569 PATTERNING OF BLOOD VESSELS | 5.620061026 |
| GO0008285 NEGATIVE REGULATION OF CELL PROLIFERATIO | 5.530971483 |
| GO0007169 TRANSMEMBRANE RECEPTOR PROTEIN TYROSINE | 5.498206806 |
| GO0007596 BLOOD COAGULATION | 5.447226092 |
| GO0008015 CIRCULATION | 5.436573095 |
| GO0017153 SODIUM DICARBOXYLATE SYMPORTER ACTIVITY | 5.391273148 |
| GO0007186 G PROTEIN COUPLED RECEPTOR PROTEIN SIGNA | 5.379357197 |
| GO0000074 REGULATION OF PROGRESSION THROUGH CELL C | 5.36917598 |
| GO0005506 IRON ION BINDING | 5.350705834 |
| GO0005529 SUGAR BINDING | 5.345748434 |
| GO0020037 HEME BINDING | 5.266458161 |
| GO0007266 RHO PROTEIN SIGNAL TRANSDUCTION | 5.239118641 |
| GO0005884 ACTIN FILAMENT | 5.198331854 |
| GO0005044 SCAVENGER RECEPTOR ACTIVITY | 5.185571699 |
| GO0008201 HEPARIN BINDING | 5.130429191 |
| GO0006835 DICARBOXYLIC ACID TRANSPORT | 5.082855262 |
| GO0030301 CHOLESTEROL TRANSPORT | 5.065979341 |
| GO0009897 EXTERNAL SIDE OF PLASMA MEMBRANE | 4.970319849 |
| GO0000302 RESPONSE TO REACTIVE OXYGEN SPECIES | 4.958212236 |
| GO0046983 PROTEIN DIMERIZATION ACTIVITY | 4.947945581 |
| GO0005540 HYALURONIC ACID BINDING | 4.868653021 |
| GO0005096 GTPASE ACTIVATOR ACTIVITY | 4.83334221 |
| GO0006979 RESPONSE TO OXIDATIVE STRESS | 4.742819776 |
| GO0007267 CELL CELL SIGNALING | 4.674093335 |
| GO0050776 REGULATION OF IMMUNE RESPONSE | 4.669760536 |
| GO0004888 TRANSMEMBRANE RECEPTOR ACTIVITY | 4.668652335 |
| GO0008305 INTEGRIN COMPLEX | 4.60079215 |
| GO0016525 NEGATIVE REGULATION OF ANGIOGENESIS | 4.531076001 |
| GO0006950 RESPONSE TO STRESS | 4.525745881 |
| GO0043123 POSITIVE REGULATION OF I KAPPAB KINASE O | 4.490558395 |
| GO0007229 INTEGRIN MEDIATED SIGNALING PATHWAY | 4.458464655 |
| GO0005792 MICROSOME | 4.45435458 |

| Gene Ontology Term | Zscore |
|---|---|
| GO0045202 SYNAPSE | -7.81405 |
| GO0042734 PRESYNAPTIC MEMBRANE | -6.05612 |
| GO0007269 NEUROTRANSMITTER SECRETION | -5.6992 |
| GO0007268 SYNAPTIC TRANSMISSION | -4.61711 |
| GO0030672 SYNAPTIC VESICLE MEMBRANE | -4.58946 |
| GO0017157 REGULATION OF EXOCYTOSIS | -4.38851 |
| GO0045211 POSTSYNAPTIC MEMBRANE | -4.22109 |
| GO0007411 AXON GUIDANCE | -4.21386 |
| GO0043005 NEURON PROJECTION | -3.79816 |
| GO0006512 UBIQUITIN CYCLE | -3.50857 |
| GO0004970 IONOTROPIC GLUTAMATE RECEPTOR ACTIVITY | -3.38013 |
| GO0005234 EXTRACELLULAR GLUTAMATE GATED ION CHANNE | -3.38013 |
| GO0006376 MRNA SPLICE SITE SELECTION | -3.18873 |
| GO0004221 UBIQUITIN THIOLESTERASE ACTIVITY | -3.16451 |
| GO0016568 CHROMATIN MODIFICATION | -2.97179 |
| GO0007270 NERVE NERVE SYNAPTIC TRANSMISSION | -2.80252 |
| GO0005832 CHAPERONIN CONTAINING T COMPLEX | -2.58938 |
| GO0016363 NUCLEAR MATRIX | -2.37003 |
| GO0035097 HISTONE METHYLTRANSFERASE COMPLEX | -2.18684 |
| GO0018024 HISTONE LYSINE N METHYLTRANSFERASE ACTIV | -2.15061 |
| GO0001675 ACROSOME FORMATION | -2.1119 |
| GO0006754 ATP BIOSYNTHETIC PROCESS | -2.05276 |

Top 100 upregulated and downregulated GO terms in CS patient cerebellum samples. An absolute value pathway Z-score cut-off of 2.0 and *p*-values ≤0.05 were considered significantly changed.

### Supplementary Table 3

The list of CS patient cerebellum samples that were used for western blot analysis.

| | UMBN | Age | Sex | Cause of death | PMI |
|---|---|---|---|---|---|
| #Ctrl-1 | 5976 | 4 | Female | Smoke Inhalation | 21 |
| #Ctrl-2 | 5928 | 27 | Female | Cardiac arrythmia | 26 |
| #Ctrl-3 | 1078 | 17 | Female | Accident, Multiple Injuries | 12 |
| #Ctrl-4 | 5391 | 8 | Male | Drowning | 12 |
| #CS-1 | 1920 | 7 | Male | Diffused alveolar damage / acute pneumonia | 16 |
| #CS-2 | 786 | 17 | female | Complications of Disorder | 22 |
| #CS-3 | 5492 | 7 | Male | Complications of Disorder | unknown |
| #CS-4 | 1286 | 5 | Male | Complications of Disorder | 14 |
| #CS-5 | 1124 | 4 | Female | Complications of Disorder | 3 |
| #CS-6 | 5105 | 8 | Male | Complications of Disorder | 6 |
| #CS-7 | 1762 | 4 | Female | Complications of Disorder | 8 |

The list of CS patient cerebellum samples that were used for western blot analysis.

### Supplementary Figure 1

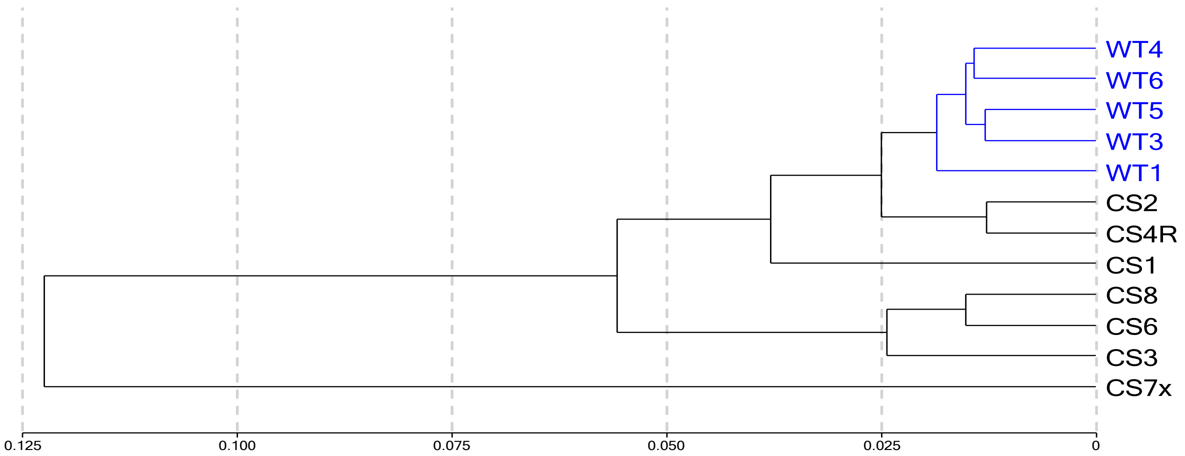

### Supplementary Figure 2

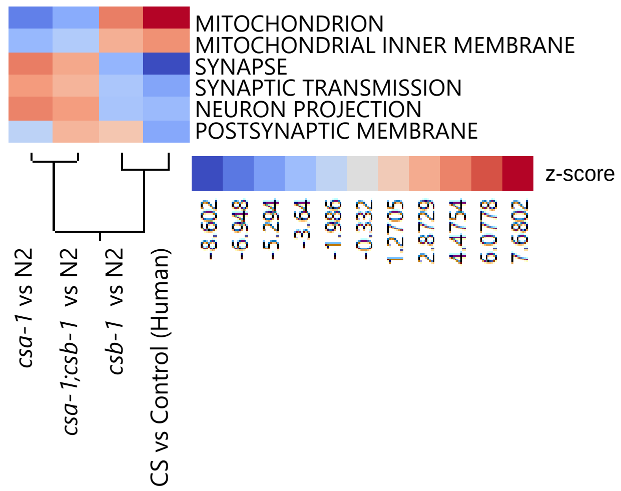

### Supplementary Figure 3

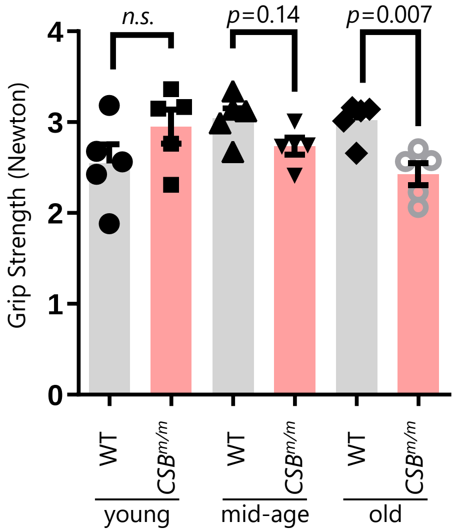

### Supplementary Figure 4

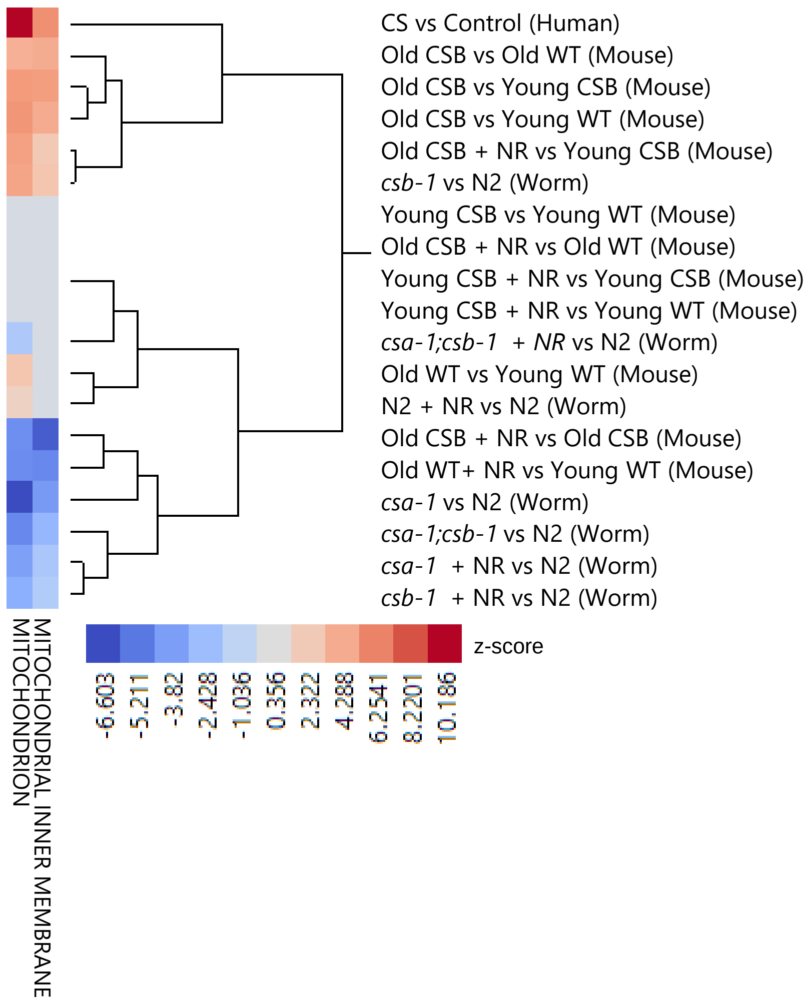

### Supplementary Figure 5

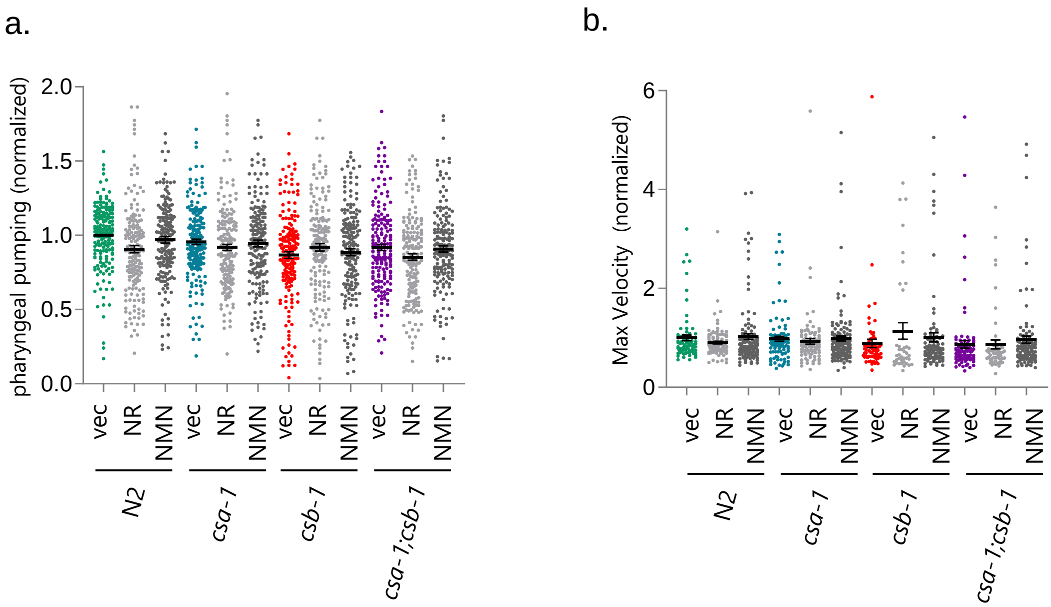

### Supplementary Figure 6

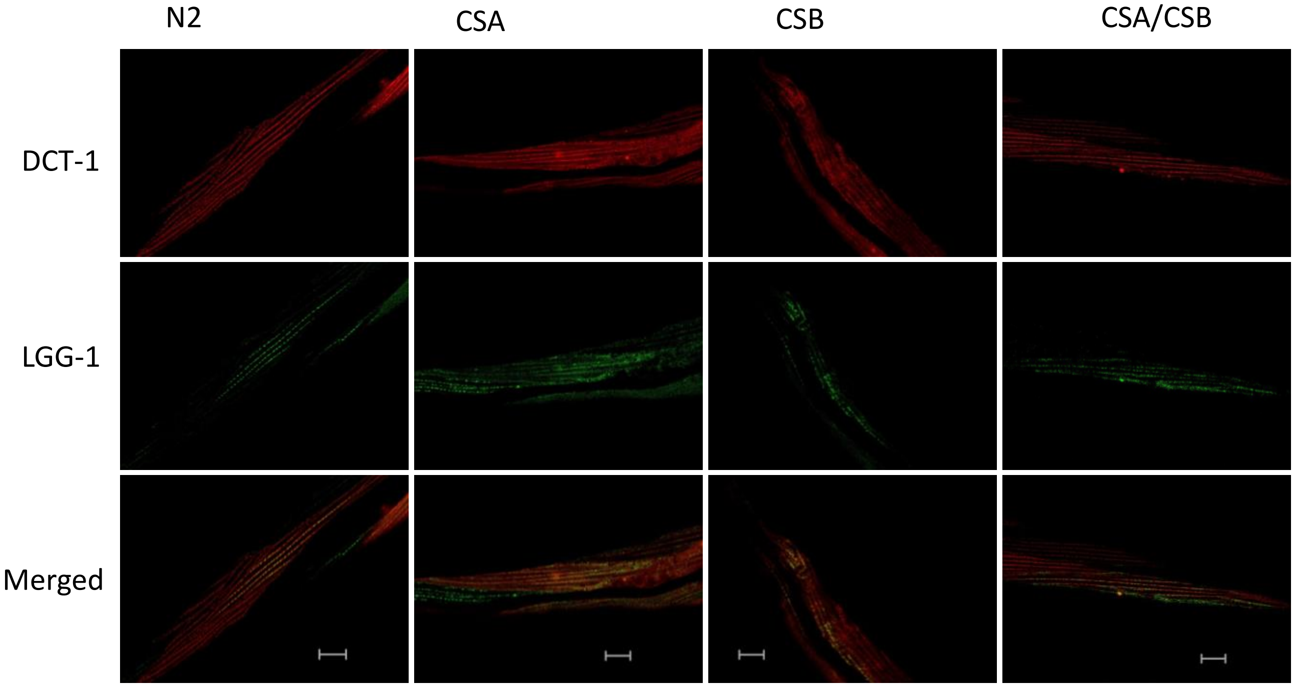
